# Ants integrate proprioception, visual context and efference copies to make robust predictions

**DOI:** 10.1101/2023.03.29.534571

**Authors:** Océane Dauzere-Peres, Antoine Wystrach

## Abstract

Feedforward models are mechanisms enabling an agent to predict the sensory outcomes of its actions. It can be implemented in the nervous system in the form of efference copies, which are copies of motor signals that are subtracted from the sensory stimulation actually detected, literally cancelling the perceptual outcome of the predicted action. In insects, efference copies are known to modulate optic flow detection for flight control in fruit flies. Much less is known, however, about possible feedforward control in other insects. Here we investigated whether feedforward control occurs in the detection of horizontal optic flow in walking ants, and how the latter is integrated to modulate their locomotion. We mounted *Cataglyphis velox* ants within a virtual reality set-up, allowing us to manipulate the relationship between the ant’s movements and the optic flow it perceives. Results show that ants do compute a prediction error by making the difference between the expected optic flow according to their own movements and the one it perceived. Interestingly, this prediction does not control locomotion directly, but modulates the ant’s intrinsic oscillator, which produces continuous alternations between right and left turns. What’s more, we show that the prediction also involves proprioceptive feedback, and is additionally modulated by the visual structure of the surrounding panorama in a functional way. Finally, prediction errors stemming from both eyes are integrated before modulating the oscillator, providing redundancy and robustness to the system. Overall, our study reveals that ants compute robust predictions of the optic flow they should receive using a distributed mechanism integrating feedforwards, feedbacks as well as innate information about the structure of the world, that control their locomotion through oscillations.

## INTRODUCTION

In 1950 von Holst and Mittelstaedt as well as Sperry found out that surgically rotating the eyes of fish or flies induced the animal to display continuous body rotations in the same direction (von Holst & Mittelstaedt, 1950, (English translation in Gallistel, 1982, Chapter 7), Sperry, 1950). To account for these results, they came up with the same concept named, respectively, efference copies and corollary discharge. These are copies of the signal sent to the motor centers that feedback to inhibit the detection of sensory information in order to subtract the part due to the movement expected. They are examples of forward models which aim at predicting the future of a current system (Webb, 2004). This enables animals to distinguish exafferences, which are external sensory stimulations, from reafferences, which are self-induced sensory stimulations due to one’s own actions. Efference copies have been demonstrated both behaviorally and neurobiologically in different animals for different contexts (Poulet & Hedwig, 2007). For example, male crickets use them to differentiate their own chirps from the chirps of other males (Poulet & Hedwig, 2006), and electric fish are effectively suppressing the sensory input that should result from their own electrical production (Kennedy et al., 2014). The same mechanisms are also responsible for the suppression of the perception of motion during saccadic eye movements in humans, so that we still perceive the world as stationary even during those saccades (Olveczky et al., 2003). In contrast, the perceived world appears to moves when pressing our eye gently with a finger, due to the lack of efference copy in this situation.

In the context of the visuomotor control of locomotion, rotational optic flow, which is defined as the perceived visual rotational movements of the scene around the observer, is important to stabilize one’s course during navigation. It has been shown that fruit flies can predict the amount of optic flow they should receive according to their own self-generated movements (Kim et al., 2015, Fenk et al., 2021). Using electrophysiological recordings, these studies demonstrated that visual cells responding to horizontal optic flow (the horizontal system north cells) are effectively inhibited proportionally to the body rotation produced by the fly. As a result, these cells respond to mismatches between the optic flow predicted and the optic flow actually detected, a so-called ‘prediction error’. Nevertheless, the existence such forward models in other insect species as well as whether proprioceptive feedback is also used to compute the prediction error and the impact of the visual context on the computation of these predicted optic flow remains largely unknown (Webb, 2004).

Horizontal optic flow is generated during locomotion when the individual is turning. Many insects, including ants (Clement et al., 2023, Graham & Collett, 2002, Lent et al., 2013), perform such turns continuously while moving, through the display of lateral oscillations, or zig-zagging (Wolf et al., 2017, Wystrach et al., 2016, Stürzl et al., 2016, Namiki et al., 2014). These oscillations are produced intrinsically by a neural oscillator located in pre-motor areas (Zorović & Hedwig 2013, Iwano et al., 2010, Namiki et al., 2014, Namiki & Kanzaki, 2016, Steinbeck et al., 2020). The horizontal optic flow generated by such oscillatory movements appears to regulate their amplitude (Clement et al., 2023, Pansopha et al., 2014, Kim et al., 2015). But what mechanisms underlie this regulation, and whether efference copies or other mechanisms are involved, remain entirely unknown.

So far, no demonstration of the existence of visual efference copies has been found in ants. Here we tackled this question with *Cataglyphis velox* ants. These solitarily foraging desert ants are expert navigators that do not use pheromone trails but rely strongly on vision for guidance (Cerdá & Retana 1997, Freas & Spetch, 2019, Mangan & Webb, 2012, Schwarz et al., 2020, Wystrach et al., 2020), making them good candidates for using visual predictions. We used a virtual-reality set-up to decouple the ants’ movement from the optic flow it received, and recorded the ants’ responses in various altered visuo-motor situations. Results reveal that ants form robust predictions of the optic flow they should expect by integrating feedforwards, feedbacks as well as innate information about the structure of the world.

## RESULTS AND DISCUSSION

### Ants’ intrinsic oscillations are modulated by the optic flow perceived

We first investigated how ants responded to optic flow by varying the visuo-motor relationship in our VR system. When in the dark, ants displayed regular oscillations between left and right turns (Fig.1A), revealing the presence of an internal oscillator, even in the absence of visual stimuli, generating those oscillations at a frequency of approximately 0.3 Hz (Mean ± SE = 0.288 ± 0,021 Hz, Fig.S1). When ants were subjected to a panorama moving with a positive gain, and thus receiving self-generated optic flow, oscillations were still present (Fig.1A). However, changing the gain from 1 (the natural visuo-motor relationship) to 3 and 5 led the ant to increase the frequency of their oscillations (LMM : 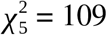, *P* < 0.001, Fig.S1) and turn at significantly slower angular speeds (LMM, 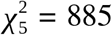, *P* < 0.001, Fig.1B). This demonstrates that optic flow impacts the expression of these oscillations, with stronger visual feedback inducing the ants to stop their current turn and turn in the opposite direction instead.

**Figure 1:**
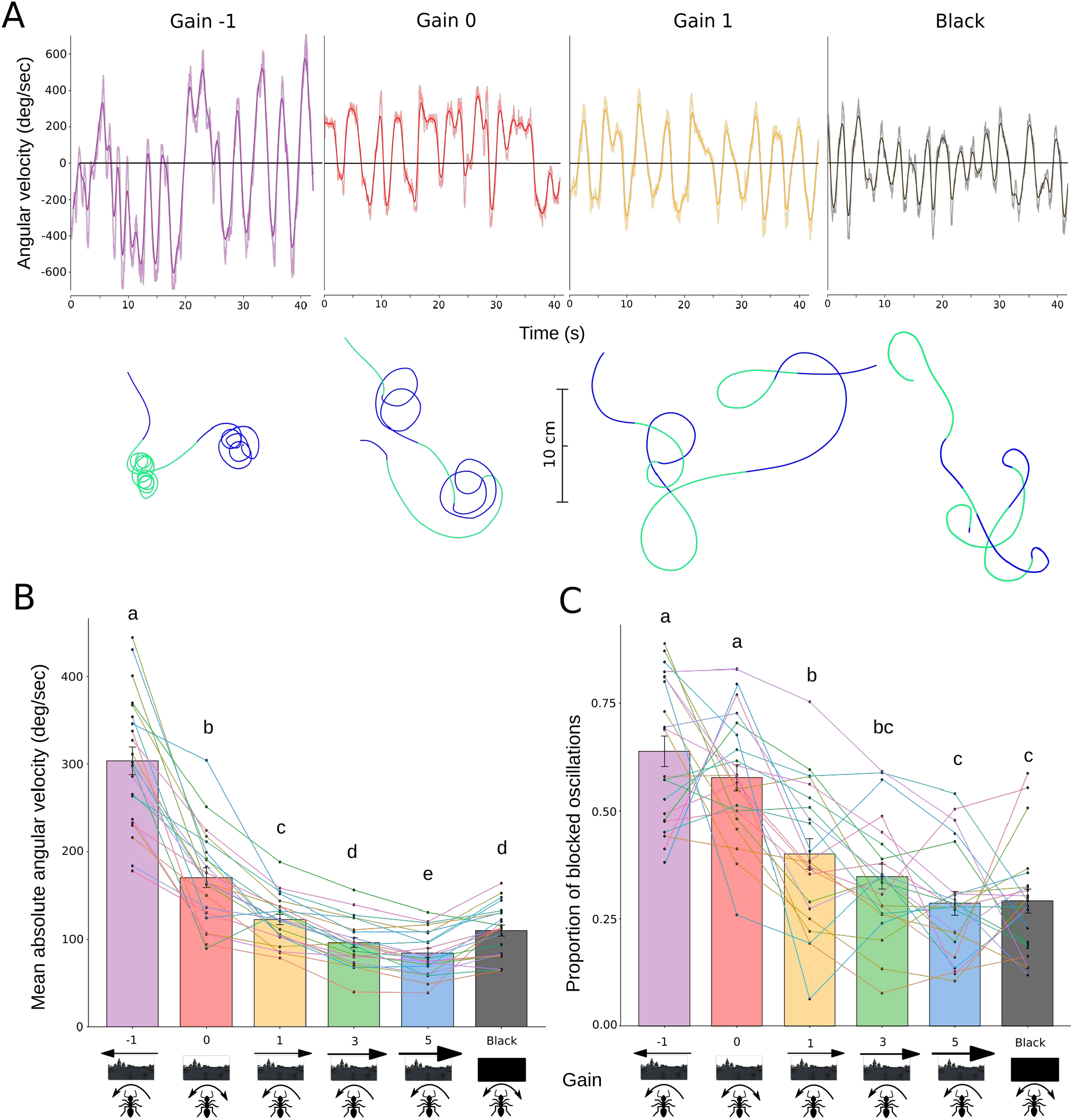
Ants with a gain of 0 behave similarly than when they are with a negative gain. **(A)** Evolution of the angular velocity signal and the associated trajectories for an individual ant between gain -1, gain 0 gain 1 and when surrounded by black. Lighter curves correspond to the raw signal while the darker ones correspond to the smoothed signal. Trajectories shown are extracts lasting 12 seconds, the blue parts correspond to right turns whereas the green parts correspond to left turns. (B-C) Mean absolute angular velocity (B) and proportion of ants’ oscillations blocked above or under 0 deg/sec (without changing direction) of ants in gains -1, 0, 1, 3, 5 and surrounded by black. Data based on 22 ants tested in the VR with both eyes uncovered. Pairs of groups that do not share a letter show significant differences in post-hoc comparisons; see statistical analysis section. The error bars shown in the bar plots (B,C) represent the standard error of the means. Each point corresponds to the response of an individual ant while the lines connect the responses of the same ant across the different conditions.

Interestingly, with a gain of –1, that is, with the world rotating in the opposite (i.e., wrong) direction in response to the ants’ movements, ants often got stuck turning in the same direction for abnormally long periods of time spanning over several oscillation cycles (Fig.1A). Interestingly, regular oscillations in angular velocity still persisted, but were shifted towards positive or negative values, meaning that ants regularly alternated between turning fast and slow in the same direction (Fig.1A, gain –1), and that the intrinsic oscillator was still influencing their behavior. There was also a significant effect of the gain on the proportion of “blocked” oscillations of the ant (defined as an oscillation in angular speed without a reversal in turning direction) (GLMM for proportional data: 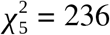, *P* < 0.001, Post-hoc statistics Fig.1C). This reveals that optic flow in the wrong direction, led to a prolongation of the current turn and/or an inhibition of the opposite turn. In short, the faster ants turn in one direction the stronger the world rotates in the wrong direction, thus motivating the ants to continue to turn in that direction, and hence getting stuck turning continuously in one direction.

### Ants compute predictions of the optic flow expected, which are silenced in the dark

We tested if ants computed prediction errors by subjecting them to a gain of 0 that is, with a static panorama. Indeed, if the optic flow detected directly controls the oscillations, the oscillator should run freely without any external influence in the absence of optic flow. Alternatively, if ants are forming predictions, the oscillator should run freely only when there is no prediction error, that is, when the perceived optic flow corresponds to the one predicted by the ants’ movements (when ants are tested with a positive gain). As a corollary, given the subtraction of the predicted optic flow, a gain of 0 (static panorama) should produce similar behaviors as a negative gain.

Remarkably, even though they received no optic flow in this condition, ants tested with a gain of 0 (static panorama) turned in a same direction for prolonged periods of time (Fig.1A), showing as much blocked oscillations as with a negative gain of –1 and significantly more than with a positive gain of 1 (or above) (Post-hoc statistics Fig.1C). This demonstrates that their oscillations are not modulated by the perceived optic flow, but by the expected optic flow; in other words, ants form a prediction error and use it to control their oscillations.

Interestingly, contrary to ants exposed to a static panorama in gain 0, ants in the dark displayed regular oscillations, even though there was no optic flow in both cases. The frequency (Fig.S1), average absolute angular velocity and proportion of blocked oscillations in the dark were significantly closer to the ones in gains 1 and 3, compared to the ones in gain 0 (Post-hoc statistics Fig.1B-C, Fig.S1). This reveals that ants must somehow silence the efference copy when in the dark so as to not compute a prediction error.

A direct prediction of those conclusions is that, at the individual level, ants rotating faster should expect more optic flow, and thus form bigger prediction errors when they are exposed to a static panorama (gain 0). On the other hand, in the dark, regardless of its angular speed an ant should not compute any prediction error. Indeed, the proportion of blocked oscillations, which is a direct indicator of prediction errors, significantly increased with the ant’s average angular speed when tested with a gain 0, but not in the dark (GLMM for proportional data: interaction, 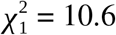, *P* = 0.001, post-hoc statistics Fig.2), the latter showing that this correlation is not simply caused by a biophysical or mathematical relationship between the two measurements (i.e., ‘angular velocity’ and ‘proportion of blocked oscillations’).

**Figure 2:**
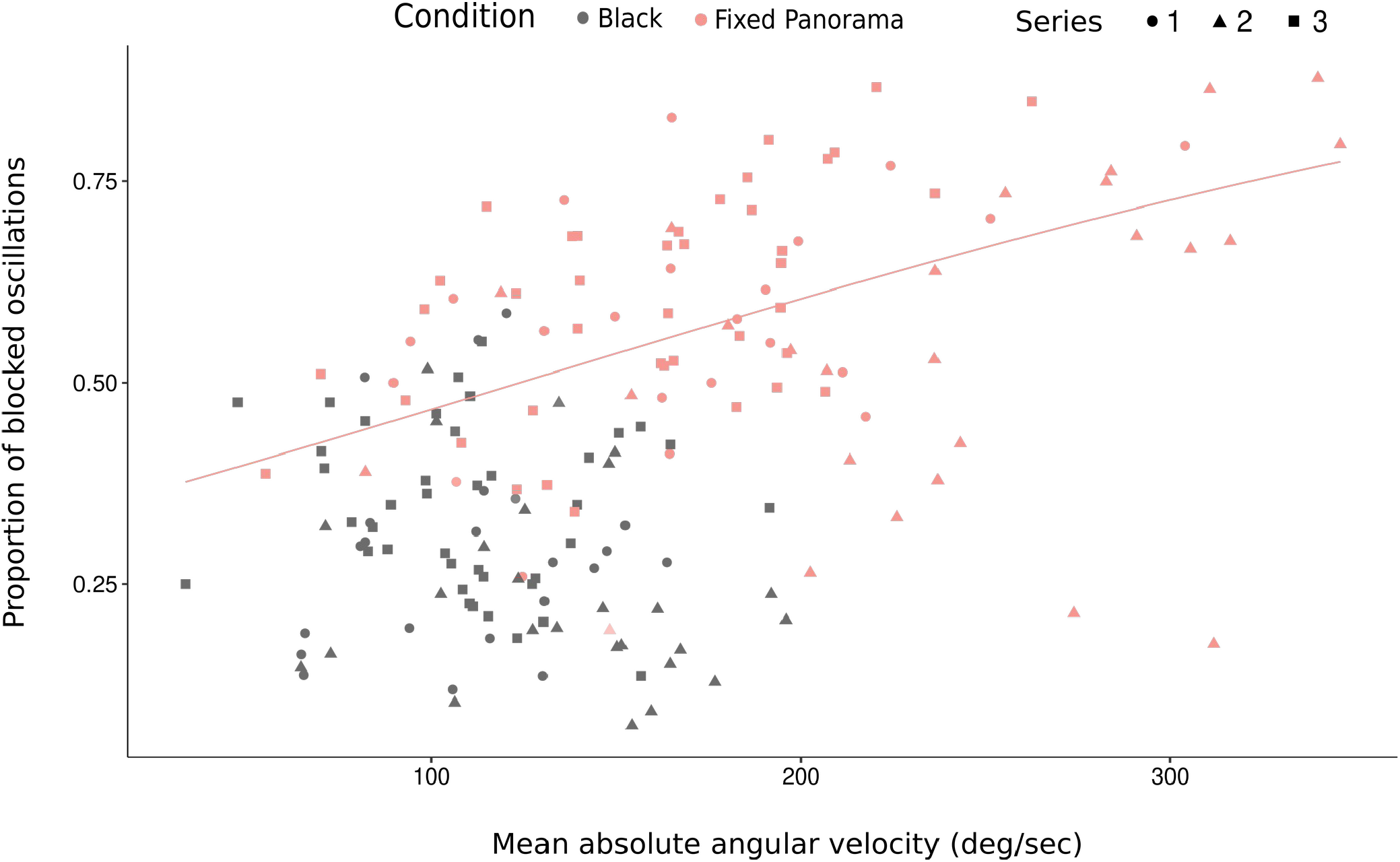
Ants turning faster get more blocked in turning in a same direction when exposed to a fixed panorama. Proportion of blocked oscillations of ants depending on their mean absolute angular velocity and the visual context they were exposed to. Pooled data based on 22 ants from the gain alteration series with both eyes uncovered (series 1), 24 ants from the visual structure alteration series (series 2) and 24 ants from the weight of the ball alteration series (series 3). There was a positive significant correlation only when the ants were exposed to a static panorama (red curve corresponds to the regression line y=exp(0.005552^*^x-0.688517)/(1+exp(0.005552^*^x-0.688517)), but not when ants where surrounded by black.

Functionally, the capacity to silence the prediction while surrounded by black seems ecologically relevant: it would certainly be disadvantageous for ants to expect to receive optic flow — and thus start turning continuously in one direction — when they are inside their nest or foraging in the absence of light. Perception of light thus appears as a necessary condition for the expression of the optic flow prediction.

### Modulation of the optic flow prediction depends on the visual context

Since ants do not expect to receive optic flow in the dark (previous section), we wondered what visual information is used to silence the optic flow prediction normally produced. Is it just the presence/absence of visual input, or can ants predict the optic flow expected based on the structure of the visual surrounding? To test this, we exposed ants to various static visual structures (gain = 0) that either should (a panorama or vertical bars) or should not (homogeneous white or horizontal bars) produce horizontal optic flow when the ant is rotating.

In all conditions without optic flow, ants displayed a significantly much higher proportion of blocked oscillations (a clear signature of prediction errors) than in the dark or when given a panorama rotating accordingly to the ants’ movement (GLMM for proportional data: 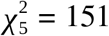, *P* < 0.001, Post-hoc statistics Fig.3B, see Fig.S2 to visualize their trajectories). This shows that in the dark, the optic flow prediction is fully canceled due to a lack of luminosity and not because the ants were able to compute the predicted optic flow by taking into account the visual structure surrounding them.

**Figure 3:**
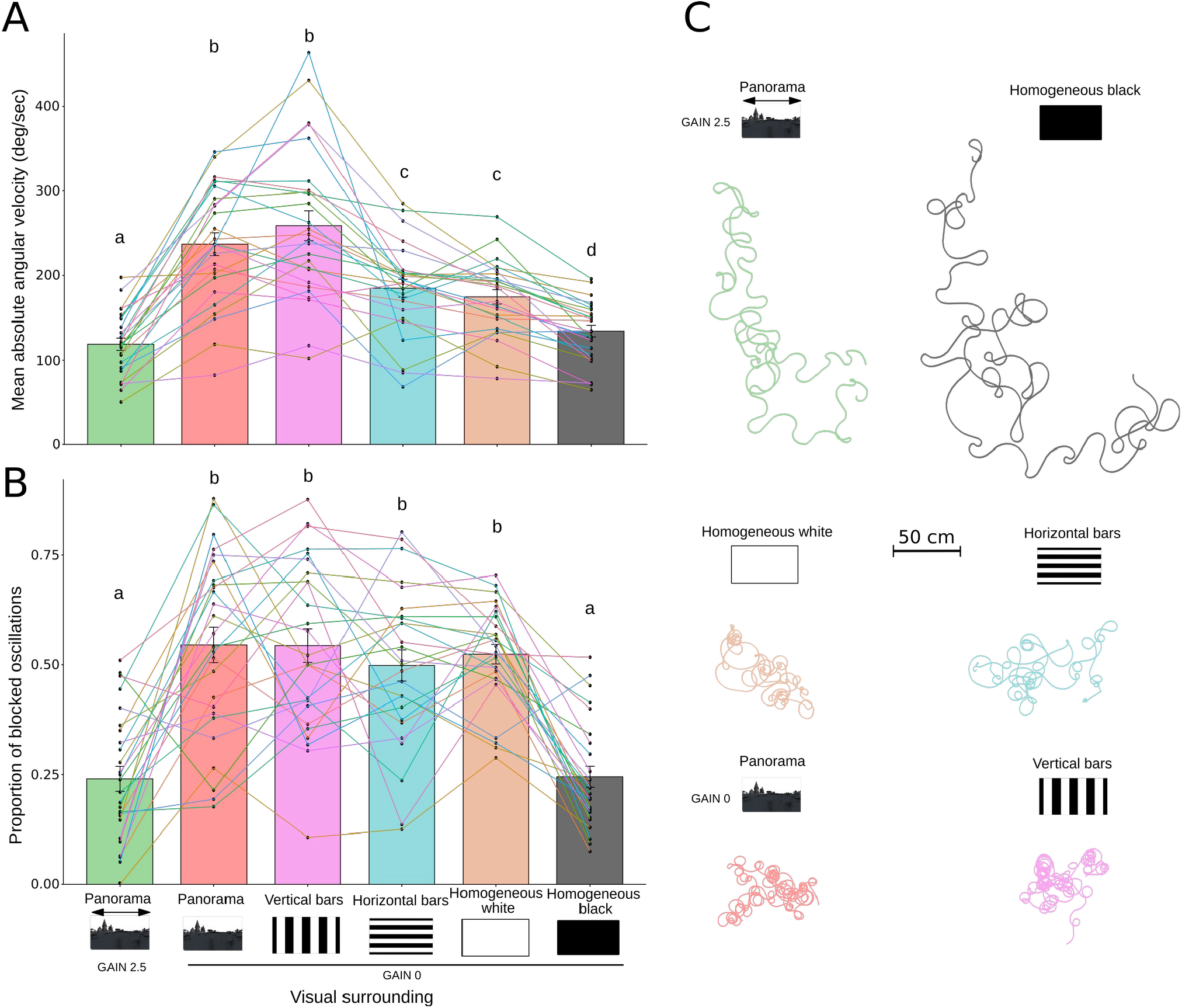
The visual structure surrounding ants impact their oscillatory behaviour. The error bars shown in the bar plots (A,B) represent the standard error of the means. Each point corresponds to the response of an individual ant while the lines connect the responses of the same ant across the different conditions. (A) Mean absolute angular velocity and (B) proportion of ants’ oscillations blocked above or under 0 deg/sec (without changing direction) of ants exposed to different visual structures in gain 0 as well as a panorama in gain 2.5. Data based on 24 ants tested in the VR. Pairs of conditions not sharing a letter show significant differences based on post-hoc comparisons; see statistical analysis section. (C) Reconstructed trajectories of an ant tested in the different visual conditions.

However, ants’ angular speed reveals that they did turn significantly quicker when facing visual structures that should (‘vertical bars’ or ‘panorama’ conditions) versus should not (‘white’ or ‘horizontal bars’ conditions) produce optic flow (LMM: 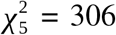, *P* < 0.001, Post-hoc statistics Fig.3A). Interestingly, this signature is analogous to what we observed between ants exposed to a panorama with a gain of –1 versus 0 (Fig.1B-C), and we know that the prediction error is greater with a gain of –1 than a gain of 0. Therefore, this suggests that the prediction error is bigger when vertical edges are present (in ‘vertical bars’ and ‘panorama’ conditions). In addition, the fact that for both parameters (mean angular velocity and proportion of blocked oscillations, Post-hoc statistics Fig.3A-B), there are no significant differences between the horizontal bars and the homogeneous white conditions, or between the panorama and the vertical bars conditions, implies that the detection of horizontal edges is clearly less important than vertical edges, if at all, for the production of optic flow predictions, and thus prediction errors.

Those results are subtle, but robust to replication (Fig.S3). What is more, a subsequent analysis showed that the number of vertical edges had an impact too, with one large vertical bar triggering lower absolute angular velocity, and thus likely less optic flow expectation, than did multiple vertical bars (Fig.S3). Overall, results show that ants do pay attention to the structure of the panorama in a functional way, with slightly higher optic flow predictions in the presence of multiple vertical edges, which indeed should produce horizontal optic flow when rotating. However, the mere presence of light (with or without horizontal edges) is already sufficient for the ant to expect optic flow, albeit less than with vertical edges.

### Efference copies or proprioceptive feedback?

The use of efference copies implies that the nature of the signal used to compute the prediction is a copy of the motor command. An alternative hypothesis, although non-mutually exclusive, is that ants could use proprioceptive feedback about the turn actually performed to derive their prediction of the optic flow that was expected. Having ants walk on a trackball allowed us to manipulate proprioceptive feedback as well as visual feedback.

Since ants mounted on the trackball need more strength to rotate a heavier ball, we manipulated the relationship between the magnitude of the motor command sent and the extent of the resulting turn, which should be faithfully detected through proprioception, by testing ants with two balls of different weights. The light ball was 1.93 times lighter than the heavy one. Thus, given the same motor strength, ants should walk and turn approximately 2 times quicker (a 93% increase) with the lighter ball (Dahmen et al., 2017). Indeed, ants’ absolute angular velocities when walking on the lighter ball significantly increased compared to when they walked on the heavier ball both in the dark and when exposed to a static panorama (LMM: 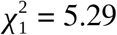, *P* = 0.021, Fig.4A). The size of this effect was not significantly different when ants were in the dark or exposed to a static panorama since there was no significant interaction of this visual context with the weight of the ball (LMM: 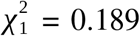, *P* = 0.664). However, this increase in speed was less than 20% (β ± SE = 17.1 ± 7,44 deg/sec), much less than the 93% increase expected. This shows that ants produced more force, and thus stronger motor signals, in the heavy ball condition, which enabled us to form the following predictions. On the one hand, if ants used proprioceptive feedback to compute their prediction, the significant increase in angular speed of the turns performed with the lighter ball means that their prediction errors (when exposed to a static panorama) should also be more important with the lighter ball. On the other hand, if ants used instead only efference copies of motor commands, the prediction errors (when exposed to a static panorama) should, in contrast, increase (or at least not decrease) with the heavier ball, given that ants produced stronger motor signal in this condition. Finally, in both cases, the weight of the ball should have no effect on the prediction error generated in the dark, since predictions are silenced in this condition (see previous sections).

**Figure 4:**
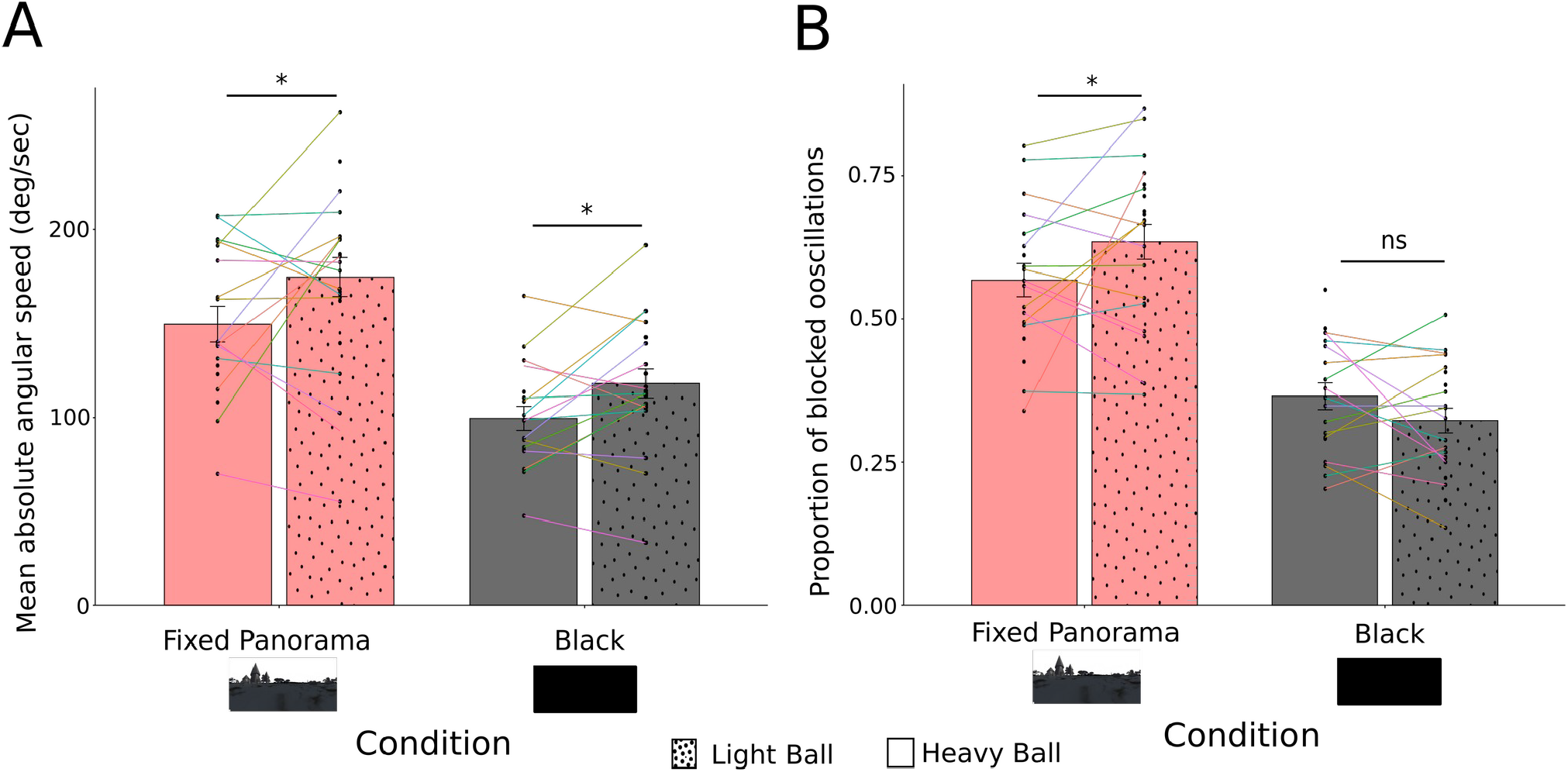
Ants walking on a lighter ball turn faster and get more blocked turning in one direction. The error bars shown in the bar plots (A,B) represent the standard error of the means. Each point corresponds to the response of an individual ant while the lines connect the responses of the same ant across the different conditions. **(A)** Mean absolute angular velocity and **(B)** proportion of ants’ oscillations blocked above or under 0 deg/sec (without changing direction) of ants walking on a heavier or lighter polystyrene ball depending on the visual context (exposed to a fixed panorama or to homogeneous black). Data based on 24 ants tested in the VR. For post-hoc comparisons : ns = non-significant,. = 0.1 < P < 0.05, ^*^ = P < 0.05, ^**^ = P < 0.01, ^***^ = P < 0.001; see statistical analysis section.

To test this, we used here again the proportion of blocked oscillations to quantify the prediction errors generated by the ants. There was a significant interaction between the weight of the ball and the visual condition on the proportion of blocked oscillations (GLMM for proportional data: 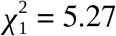, *P* = 0.022, Fig.4B). As expected, there was no significant effect of the weight of the ball in the dark. When exposed to a static panorama, however, ants displayed significantly more blocked oscillations when walking on the lighter ball (Post-hoc statistics Fig.4B). This indicates that their prediction errors, and therefore their predictions, were larger when walking on the lighter ball. Those results strongly suggest that ants are using proprioceptive feedback to generate predictions about the optic flow they expect. However, they do not reject the possibility that they are also using efference copies of motor information. In fact, other data points toward the idea that ants integrate both sources of information.

As we discussed previously, ants’ oscillations when surrounded by black are *not* modulated by optic-flow prediction errors; in other words, oscillations in the dark should be similar to when the visual panorama produces optic flow matching exactly the one expected by the ants’ movements (i.e., with a gain of 1). Interestingly, ants’ oscillations (absolute angular velocity and proportion of blocked oscillations) when surrounded by black and exposed to a panorama with a gain of 1 did not match. Instead, the ants’ behavior in the dark matched significantly best with a gain between 1 and 3, closer to 3 than to 1 (Fig.1B-C). With a gain of 1, our VR setup generated a rotation of the scenery corresponding exactly to the rotation of the ball, that is, matching the turns physically produced by the ant, so it appears surprising that an optic flow 3 times stronger would be closer to the optic flow ants expected to receive if the ant used purely proprioceptive feedback. We could, however, explain this difference if ants also used a motor efference copy. Ants mounted on the trackball need more strength to rotate the ball than they would normally to rotate their body on solid ground, around a factor of 100 to 3000 according to Dahmen et al., 2017. With an efference copy of the motor command, the expected optic flow will thus be over-estimated compared to the actual rotation of the ball. With a gain of 1, the panorama should therefore rotate slower than what the ant is predicting, leading to the mismatch observed with oscillations in the dark, and explaining why a gain more important than 1 is necessary for optic flow to match the ants’ predictions.

### An additional ‘proprioceptive-to-motor’ loop

The fact that oscillations in the dark matched best with a gain around 2.5 could result from several possibilities. Assuming, conservatively, that the ball requires at least 100 times the power needed for the ant to rotate normally (Dahmen et al., 2017), the prediction derived from a motor efference copy would match with a gain of 100 while the prediction derived from proprioceptive feedback would fit a gain of 1. The resulting gain of 2.5 could be explained by having proprioceptive feedback information much more weighted than motor efference copies when computing the prediction errors. This explanation alone, however, is unlikely. Indeed, the fact that ants on the ball in the dark do perform oscillations with amplitudes that are roughly comparable to what is observed on the ground, and definitely not 100 times smaller, shows that a regulation of the turns’ amplitude based on proprioception is also taking place independently of vision. This control loop is certainly regulating the strength actually needed to reach the desired amplitude, as informed by proprioception, and we can envision several ways how this could be implemented (Fig.5). Nevertheless, the difference in angular velocity observed with balls of different weights (Fig.4A) shows that this control loop does not compensate exactly for the difference in strength needed to reach a same amplitude. Such a gap suggests that the ants prediction still overestimates of the movement that will be effected on the ball, and hence the efference copy must also overestimates the expected optic flow, which explains why the apparent absence of prediction errors occurs for gain >1, here around 2.5.

**Figure 5:**
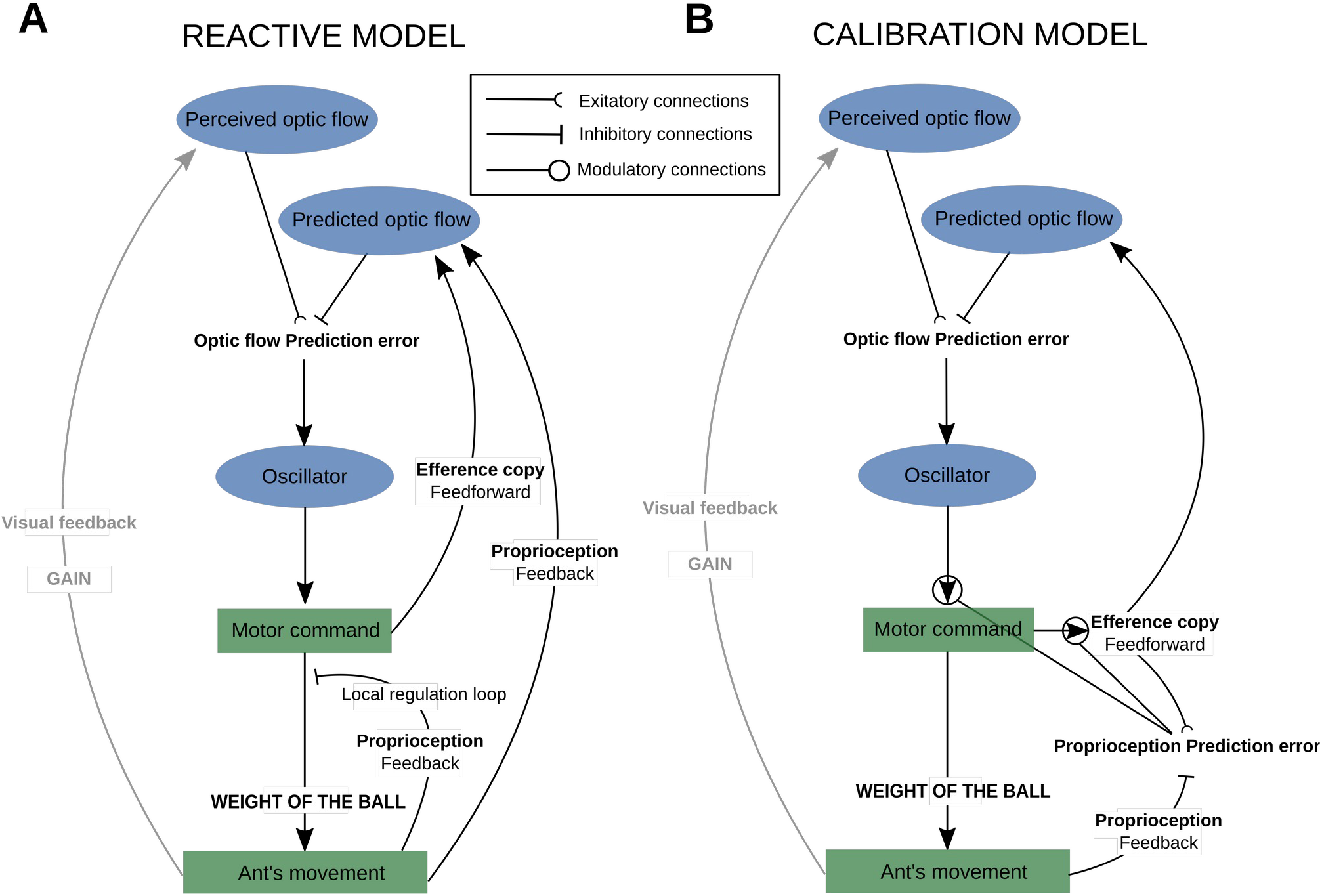
Visuo-motor control of ant’s oscillations. The prediction error is computed as the difference between the optic flow perceived and the one predicted. The perceived optic flow is a direct result of the ant’s movement multiplied by the gain we set up for our experiment (1 in natural condition). The prediction error then impacts the generation of oscillations by increasing the current turn if the individual rotated less in the correct direction than expected, and inhibiting it if the opposite is true. **(A)** Reactive model. The predicted optic flow is computed using both motor efference copy and proprioceptive feedback. A local loop, independent of vision, regulates the strength of the motor commands according to the proprioceptive feedback **(B)** Calibration model. The predicted optic flow is solely based on a motor efference copy but the gain of this efference copy is calibrated by a proprioceptive feedforward loop minimizing the proprioception prediction errors. In addition, the proprioception prediction errors also calibrate the gain of the motor command to keep the initially desired turn amplitude.

### Summation of the prediction errors

We next wondered how optic flow information from both eyes is combined before modulating the oscillations. To do so we exposed one of the ant compound eyes to the dark by covering it with paint (Fig.7C). There was a significant interaction between the gain and the number of eyes uncovered (1 or 2) on both the proportion of blocked oscillations (GLMM : 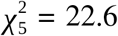, *P* < 0.001, Fig.6B) and their absolute angular velocity (LMM : 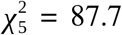, *P* < 0.001, Fig.6A). While the amplitude of monocular ants’ oscillations and their probability to get blocked were reduced in gain -1 and 0, this probability had a tendency to increase in gain 5 (Post-hoc statistics Fig.6A-B). In other words, covering one of the ant’s eyes (either the left or the right one) significantly reduced the effects of gain alteration compared to their default behavior in the dark. Gain –1 and 0 normally produce negative prediction errors leading to bigger turns than in the dark. Conversely, Gain 5 produces positive prediction errors leading to smaller turns than in the dark. In both cases, the behavioral effects of the prediction errors were reduced with one eye covered. These results show that the prediction errors stemming from both eyes are integrated before modulating the oscillator. Covering an eye simply led to no prediction error for this eye (as observed in the dark), and thus a reduced overall prediction error. Note that if the prediction error of the covered eye was ignored, or if only the most important prediction errors between the two eyes were being used, we should not observe those intermediate behaviors. This also explains why there was no significant effect of covering the ants’ eye when no prediction errors were expected (between gain 1 - 3). Therefore, under normal conditions, both eyes send similar redundant information, which contributes to the robustness of the system and makes it more accurate.

**Figure 6:**
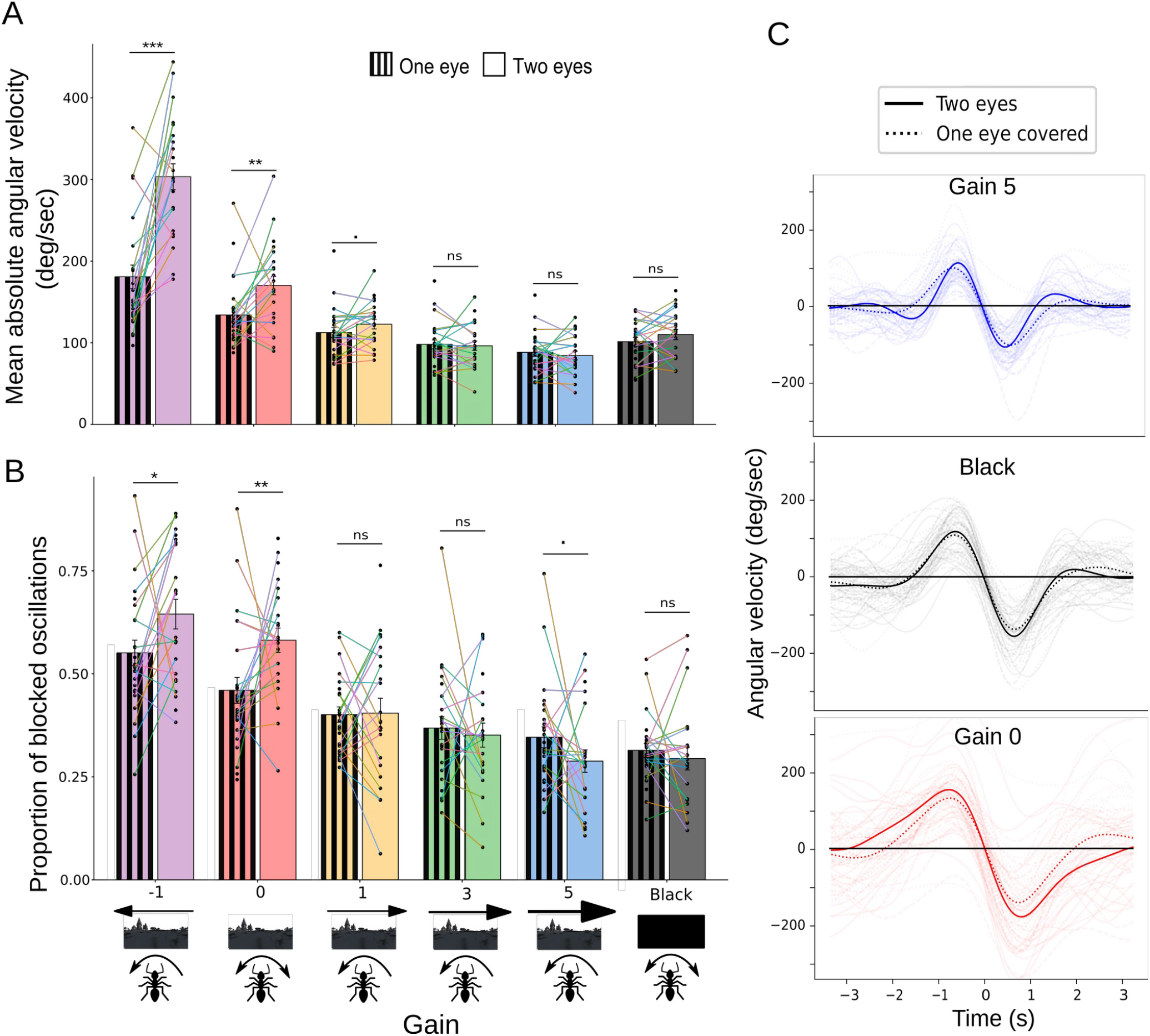
Covering one of the ants’ eye decreases the effect of gain modulation compared to when they are in the dark. The error bars shown in the bar plots (A,B) represent the standard error of the means. Each point corresponds to the responses of an individual ant while the lines connect the responses of the same ant across the different conditions. For post-hoc comparisons : ns = non-significant,. = 0.1 > P > 0.05, ^*^ = P < 0.05, ^**^ = P < 0.01, ^***^ = P < 0.001 ; see statistical analysis section. (A-B) Impact of the number of eyes uncovered on (A) the mean absolute angular velocity and (B) the proportion of oscillation blocked above or under 0 deg/sec (not changing direction) of ants depending on the gain in closed-loop. Data based on 27 ants tested in the VR including 25 with one eye covered and 22 with both eyes uncovered.(C) Mean oscillation cycles of ants with both eyes uncovered or with one eye covered in gain 0, gain 5 and surrounded by black. Curves with lighter shades correspond to the average oscillations of individual ants, while the dark ones are averaged at the population level.

**Figure 7.**
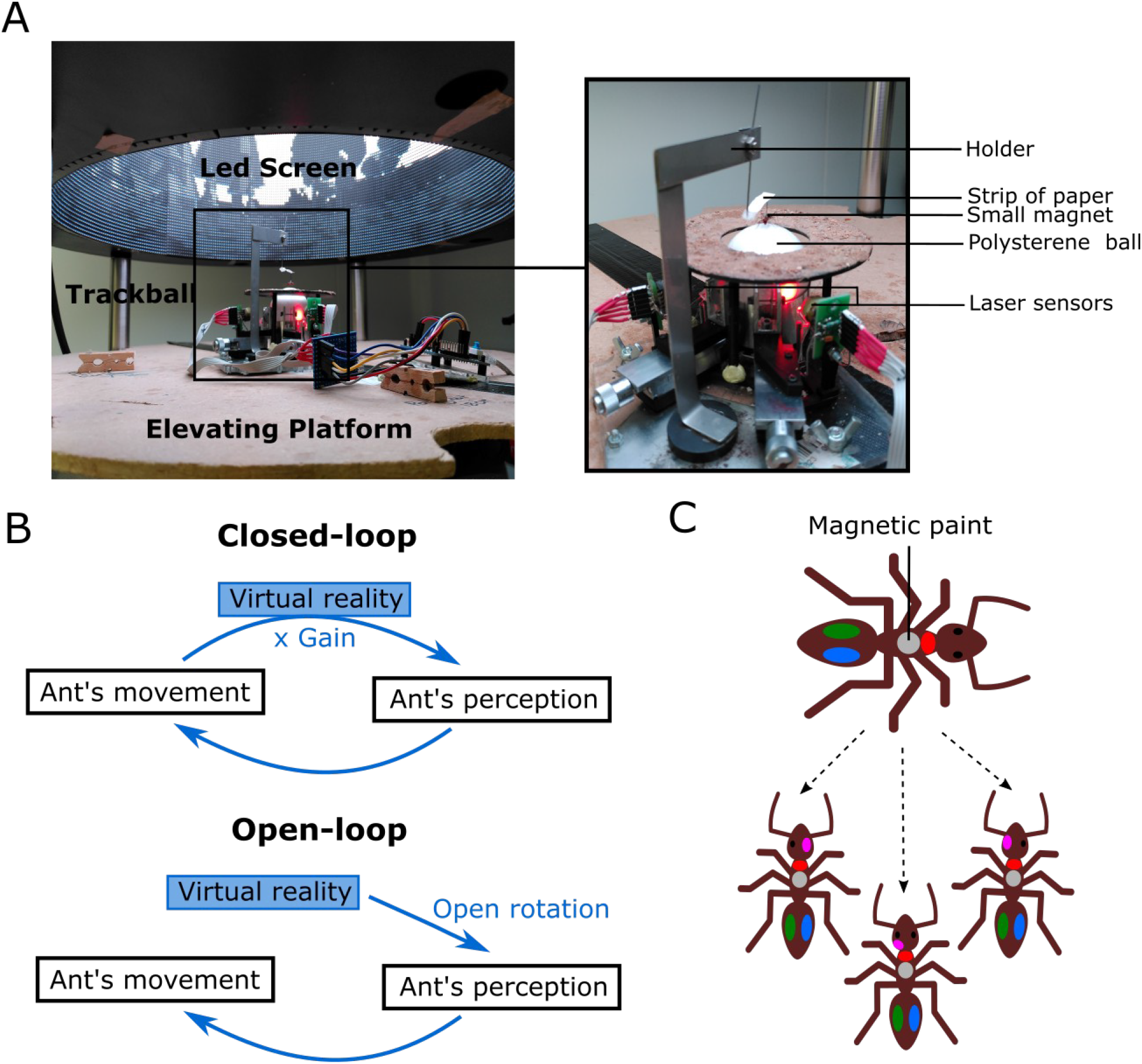
Virtual reality experimental set-up. **(A)** Pictures of the virtual reality arena and the trackball **(B)** Operating modes of the virtual reality **(C)** Placement of the paint on ants before getting tested in virtual reality.

### General Discussion

In insects, the Lateral Accessory Lobes (LAL) are pre-motor areas responsible for the generation and control of oscillations or other steering mechanisms such as phototaxis (Clement et al., 2023, Iwano et al., 2010, Namiki & Kanzaki, 2016, Namiki et al., 2014, Steinbeck et al., 2020). Some studies have highlighted the importance of optic flow in regulating the oscillations (Clement et al., 2023, Kim et al., 2015, Pansopha et al., 2014). Here we demonstrated that, at least in ants, what modulates the LAL is not directly the optic flow detected, but the prediction errors resulting from the difference between what was expected and what was perceived. Interestingly, regular oscillations persisted across all our manipulations but were expressed with various amplitudes and frequencies, showing that prediction error signals to the LAL modulate the intrinsic oscillators rather than bypass it. This corroborates the idea that such intrinsic oscillations are an ancestral feature in insects that has been outsourced for various tasks through the control by various modalities (Wolf et al., 2017, Wystrach et al., 2016, Stürzl et al., 2016, Namiki et al., 2014, Namiki & Kanzaki, 2016, Steinbeck et al., 2020). In addition, there seems to be a summation of the prediction errors from both eyes. *Cataglyphis* ants’ brains actually have a neural track, called the Inferior Optic Commissure (IOC) crossing the hemisphere to connect both left and right optic lobes. The IOC also project to the ventrolateral neuropils, which are known to possess wide-field optic flow detector cells in flies (Habenstein et al., 2020). This corroborates the assumption that the prediction errors are computed in each eyes and then summed upstream from the LAL, rather than being purely ipsilateral.

Our results highlight common points but also differences with the way flies deal with optic flow. Like for our ants, electrophysiological recording in fruit flies showed that the strength of the efference copy depended on the structure of their visual surrounding: the magnitude of the efference copy sent for the same movements was weaker when surrounded by a homogeneous grey screen compared to when they were exposed to vertical bars (Kim et al., 2015). This suggests that modulation of the predicted optic flow produced by the visual structure is ancestral to both ants and flies. However, contrary to ants in the dark, the expression of the efference copy signals in flying flies is not silenced when blinded through artificially prolonged depolarizing afterpotential (PDA) causing R1–6 opsin to photoconvert to a persistently active state (Kim et al., 2015), even if it remains doubtful that flies would get blocked turning into circle when in the dark (see for example Seelig & Jayaraman, 2015 for flies walking in the dark).

Also, we showed that optic flow prediction errors in ants are based at least in part on proprioceptive feedback, and not purely on motor efference copies. In insects, connections from proprioreceptors to leg motor neurons and their role in locomotion control has been largely demonstrated (Höltje & Hustert, 2003, Kukillaya et al., 2009, Tuthill & Azim, 2018, Tuthill & Wilson, 2016), but we show here that proprioceptive feedbacks also send connections to the brain to participate to the visual predictions. While efference copies are faster because they are produced before the movement and thus probably important for fast flight control, proprioceptive feedback signals may be more reliable and therefore weighted more when walking than flying, and thus perhaps weighted more in ants than in flies. Contrary to efference copies, proprioceptive feedback do not produce a strico-sensus “prediction” since the signal must arrive with a delay, after that the corresponding optic flow has been perceived. However, because this time lag is likely under 30 ms (Gebehart & Büschges, 2021) while an oscillation spans over several seconds, proprioceptive feedback can certainly still be used, in addition to motor efference copies, to the estimation of the currently expected optic flow (Fig.5A, Reactive model). Alternatively, proprioception could participate to the predicted optic flow indirectly, by calibrating the gain of the motor efference copy according to the mismatch (error) between the motor command effected and the proprioceptive feedback (Fig.5B, Calibration model). Note that both ‘reactive’ and ‘calibration’ models are not mutually exclusive (Fig. 5).

Finally, it is interesting to note that when exposed to a negative gain for prolonged periods of time (around 8 sessions of 4 minutes), fruit flies learned to reverse their movements in order to stabilize the panorama in front of them (Heisenberg & Wolf, 1984, Wolf et al. 1992). This may be an adaptation to flight control, where stabilizing the perceived panorama is crucial. Whether our ants are also capable of recalibrating their optic flow prediction errors across time, and how such plasticity is neurally implemented in insects remains to be seen, and would form an integrating agenda for future research.

## METHODS

### Studied species and housing conditions

All the experiments were done using *Cataglyphis velox* desert ants. Those ants can measure up to 12 mm (Bocher et al., 2007), which facilitates manipulations such as covering their eyes and putting them in virtual reality. Colonies comprising one or more queens, workers and larvae were collected in Seville (Spain) in 2020 and brought to the CRCA in Toulouse. These colonies were kept in vertical nests made by digging galleries and chambers in aerated concrete. The nests were kept in the dark in an air-conditioned room under controlled conditions (temperature 24–30°C and humidity 15–40%) and the foraging area was exposed to a 12/12h day/night luminosity cycle. Nests were connected by a transparent tube to a 40 × 30 cm foraging arena covered by sand and exposed to a heating lamp following the 12/12 cycle. In this arena workers always had access to water as well as sugar water (40% sugar for 60% water). In addition, pieces of mealworms were dropped in the arena 3 times per week.

### Experimental set-up : Virtual world and data acquisition

We used a Virtual Reality (VR) setup made up of a trackball system surrounded by a 360° cylindrical LED screen (Fig.7A). The trackball enabled a tethered ant to walk on a polystyrene ball floating on a cushion of air compensating its movement (Dahmen et al., 2017). Ants received a dot of magnetic paint on their thorax and were mounted on the trackball in a given body orientation using a holder with a micro-magnet. The trackball was placed on a lifting platform enabling to raise the mounted ants within the VR screen. The VR cylindrical screen (50cm diameter and 76cm high) was composed of 73728 RGB LEDs, offering a resolution of 0.94 deg/pixels, which is higher than *Cataglyphis* ants’ eyes resolution (>2deg/pixels, Zollikofer et al., 1995). The scenery projected on the screen was controlled online by a computer using custom codes implemented in a Unity® 2020.1 environment. The movements of the ants were recorded by two sensors pointing at the ball and were sent back to the computer to update the scenery depending on the parameters chosen. A camera placed on the top of the VR set-up recorded the ant from above and enabled us to check whether the ant was always attached properly.

In all experiments, we chose to display a black and white panorama mimicking the natural habitat of *C*.*velox* and producing realistic optic flow information. In some experiments, the screen could display uniform black or white, as well as vertical or horizontal strips. The VR enabled us to manipulate the relationship between the rotational movements performed by the ants and the visual rotation of the scenery. For all of the experiments in this study, we only focused on the rotational component and got rid of the influence of translational movements: in other words, the scenery around them could only rotate but not change. We chose to do that to ensure that all ants were exposed to horizontal optic flow of the same magnitude.

In some conditions, the VR system was set in ‘closed-loop’: the ants’ movement affected the movement of the panorama on the screen, and we could manipulate this relationship through a parameter called “gain” (Fig.7B). A gain of 1 means that the rotation of the scenery on the screen matches exactly with the ball’s rotational movement produced by the ants, mimicking what happens in the real world. A gain of 5 means that the scenery rotates 5 times faster; and a gain of –1 means that the scenery rotates at the correct speed but in the opposite direction to what it normally should according to the ant’s movements. In other conditions, the VR system was set in ‘open-loop’: the ants were exposed to a fixed image with a chosen structure, and their movements had no impact on the visual feedback, which is equivalent to a gain of 0.

### Experimental protocol

We conducted several series of experiments. In all experiments, tested ants were randomly picked from the foraging area and marked individually using a color code composed of a dot of paint on their thorax and two dots on their abdomen. Magnetic pain was also applied on the middle of their thorax to enable to mount them on the trackball (Fig.7C).

The painted ants were then placed for 2 min in a small pot to let the paint dry, and were mounted on the trackball with the platform on the lowest position. The screen of the VR was first turned black while the platform was raised to immerse the ant in the VR setup; the screen was then switched on and the trackball movements were recorded. Once the trial was over the ant was placed back in its nest.

For the first series of experiments, another dot of paint was either applied on their left eye (LE), right eye (RE), or above their left eye (2E) for the sham group (Fig.7C). At the end of the trial, the painted eye cap was then removed before releasing the ant back to its nest. Individually marked ants could be picked up from the nest multiple times to be tested in the different conditions (LE, RE and 2E) in a randomized order. In total 27 different ants were tested in closed-loop under six conditions successively without leaving the VR: they were tested with a gain equal to –1, 0, 1, 3, 5 and with the screen filled with black, in a pseudo-randomized order. Each condition lasted for 90 seconds and the screen was filled with black for 15 seconds between two conditions. When the gain was equal to 0, to prevent any bias across ants, the fixed panorama was rotated so that all the ants perceived it from the same orientation, whatever their absolute orientation on the trackball. To do so, the ant’s body orientation was measured using the camera, and the panorama orientation was adjusted during the 15 seconds of darkness preceding the gain 0 condition.

For the second series of experiments, 24 different ants (which were not used in the first series) were tested successively without leaving the VR in 6 visual conditions: one in closed-loop with a gain of 2.5 and a realistic panorama, and five in open-loop (i.e., a gain of 0) with different visual structures: horizontal bars, vertical bars, homogeneous white, homogeneous black, and the realistic panorama. The ants were exposed to all the conditions in a pseudo-randomized order. In the horizontal- and vertical-bars conditions we controlled for the overall luminosity by ensuring that the proportion of black vs. white LEDs remained constant and equal.

For the third series of experiments ants were first tested in the VR randomly with a lighter polystyrene ball of 229 mg or with a heavier polystyrene ball of 442 mg. Individually marked ants could be picked up again from the nest latter to be tested with the other ball they were not first tested with. In total, 24 different ants (not used in the first two series) were exposed, for each ball condition, to the dark and to a fixed panorama (i.e. gain 0) successively without leaving the VR in a randomized order. Each visual context condition lasted for 90 seconds and the screen was filled with black for 15 seconds between the two.

### Data transformation

All statistical analyses were done using R v. 4.0.3 (R Core Team, 2020) whereas the transformation of the raw data acquired by the system into exploitable variables and their visualization was done with Python v. 3.9.7.

For all the trials of each ant, the raw data collected were split between the different tested conditions and transformed into understandable measures thanks to a python code. That way, the virtual trajectory as well as the angular velocity over time (negative when going right and positive when going left) were extracted. To decrease the natural noise of the recording system (recording at Mean ± SD = 45.3 ± 1.88 fps), the raw angular velocity was resampled to have a data point every 33 μs (30 fps) and smoothed by running twice an average filter with a sliding window of 20 frames long.

### Statistical analysis

Two response variables were computed for each condition in every trial for all ants: the average absolute angular velocity (corresponding to how quickly an ant turned no matter the direction) and the proportion of detected oscillations that were defined as “blocked” because they didn’t involve switching direction (characterized by not crossing 0 deg/sec).

Analyses were done using linear mixed-effect models (LMMs) or generalized linear mixed-effect models (GLMMs) based on restricted maximum likelihood estimates with the R package lme4 (Bates et al., 2015). In all the models, the identity of the ant was included as a random factor to account for repeated measurements, as well as the identity of the trial when ants were tested in the VR more than once. For all the models, the normal distribution of residuals as well as the homogeneity of variances was verified by visually checking, respectively, QQplots and the residuals versus the fitted values (Faraway, 2006). For the GLMMs we checked for overdispersion using the function check_overdispersion of the R package performance (Lüdecke et al., 2021); if overdispersion was detected we added observation-level random effects as a way to deal with this overdispersion (Harrison, 2014). P-values were calculated by type II or type III (if interactions were included and significant) Wald chi-square tests. Pairwise post-hoc comparisons on significant (G)LMMs were done by using the same model on subsets of the data including only the groups to compare to test our hypotheses. When a given group was involved in more than one such post-hoc comparison, The p-values obtained (with the same test as the original model) were corrected using the sequential Bonferroni correction after Holm according to the comparisons we were interested in, and compared to a significance level of 0.05 (Holm, 1979).

We decided to regroup the two conditions “right eye covered” and “left eye covered” into a single condition “one eye covered”, after checking there was no significant difference between covering the right or the left eye on the response variables tested (see Fig.S4). The values associated with the resulting condition “one eye” were obtained by calculating for each ant the average between their values with the left eye covered and the right eye covered; if the ant only got tested with one of the two possible eyes covered, the value associated was kept for the “one eye” condition. To facilitate the understanding of the study, the results of those models will be split up: the comparisons between the different groups will be reported in different corresponding sections.

#### Series 1: Alteration of the Gain

We tested for the effect of the gain (factor with 6 levels: –1, 0, 1, 3, 5 and surrounded by black) and the number of eyes uncovered (factor with 2 levels: 1 eye or 2 eyes) as well as the interaction between the two factors on the proportion of ants’ oscillations that were “blocked” in one direction and their average absolute angular velocity. We used a LMM for the absolute angular velocities with the log transformation, and a GLMM for proportional data with the binomial distribution and the link function logit for the proportion of blocked oscillations. Since the interaction was significant for both models, we chose to look at all the pairwise post-hoc comparisons between the groups with 2 eyes and at the comparisons between 1 versus 2 eyes inside each gain group.

#### Series 2 : Alteration of the visual structure

We tested for the effect of the type of visual structure ants were exposed to (factor with 6 levels: fixed horizontal bars, verticals bars, homogeneous white, homogeneous black, static panorama and a panorama in closed loop (gain of 2.5)) on their average absolute angular velocities and the proportion of their oscillations which is blocked. We used a LMM for the absolute angular velocities with the log transformation, and a GLMM for proportional data with the binomial distribution and the link function logit for the proportion of blocked oscillations.

#### Series 3 : Alteration of the weight of the ball

We tested for the effects of the weight of the polystyrene ball (factor with 2 levels: heavy or light) and the visual context (factor with 2 levels: in the dark or exposed to a fixed panorama) as well as the interaction between the two factors on ants’ average absolute angular velocities and the proportion of their oscillations which was blocked. We used a LMM for the absolute angular velocities, and a GLMM for proportional data with the binomial distribution and the link function logit for the proportion of blocked oscillations. Since the interaction was significant for both models, we chose to look at the pairwise post-hoc comparisons between light versus heavy ball inside both visual context groups.

#### Series 1, 2 and 3

We pooled together the data of all the series where ants were exposed to the dark and to a static panorama (gain 0). We then tested the correlation between their proportion of blocked oscillations (response variable) and the average absolute angular velocity (predictor variable) by taking into account the visual context (factor with 2 levels: surrounded by black or surrounded by a static panorama) and the intectaion between the two factors with a GLMM for proportional data with the binomial distribution and the link function logit. In addition to the identity of the ant and the identity of the trial, we added the identity of the series as a random factor to account for the similarity in the context of the measure (other conditions tested). Since the interaction between the visual context and the average absolute angular velocity was significant we carried out post-hoc analysis by testing the effect of the average absolute angular velocity separately on both groups.

## ACKNOWLEDGMENTS

We want to thank Sebastian Swartz for maintaining the colonies as well as his help in teaching how to manipulate ants and Ken Cheng for his helpful comments on the manuscript.

## ADDITIONNAL INFORMATION

### Funding

Funder: European Research Council,

Grant reference number: EMERG-ANT 759817,

Author: Antoine Wystrach

### Author contributions

ODP Conception and Design of experiment, Data collection, Analysis and interpretation of data, Drafting and revising the article. AW Conception and Design of experiment, Analysis and interpretation of data, Drafting and revising the article, Supervision of project.

## SUPPLEMENTARY INFORMATION 1

**Figure S1:**
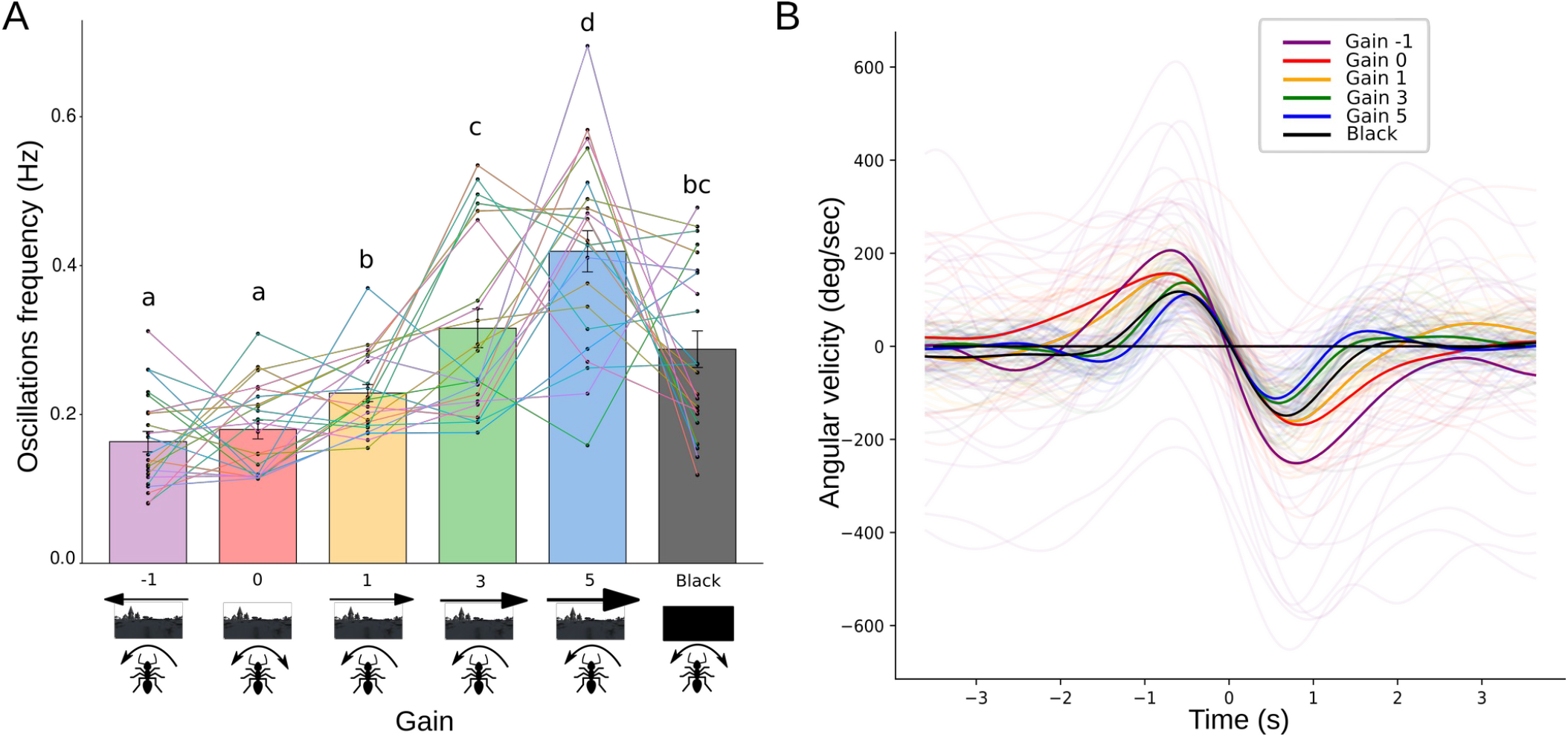
Ants alternate more frequently between left and right turns when the world moves faster than them. **(A)** Frequency of the oscillations of ants in gains -1, 0, 1, 3, 5 and surrounded by black. Data based on 22 ants tested in the VR with both eyes uncovered. Frequencies were obtained by using Fourier transform on the smoothed angular velocities signals. Different letters correspond to significant post-hoc comparisons; see the statistical analysis section. The error bars represent the standard error of the means. Each point corresponds to the response of an individual ant while the lines are connecting the responses of the same ant across the different conditions. We used a LMM to test the effect of the gain on the frequency of ants’ oscillations. The identity of the ant was added as a random factor to account for repeated measurements. There was a significant effect of gain on the oscillations frequency (LMM : 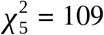, *P* < 0.001). Different letters correspond to significant post-hoc comparisons using the sequential Bonferroni correction after Holm. **(B)** Mean oscillation cycles of ants with both eyes uncovered in gains -1, 0, 1, 3, 5 and surrounded by black. Curves with lighter shades correspond to the average oscillations of individual ants, while the dark ones are averaged at the population level.

## SUPPLEMENTARY INFORMATION 2

**Figure S2:**
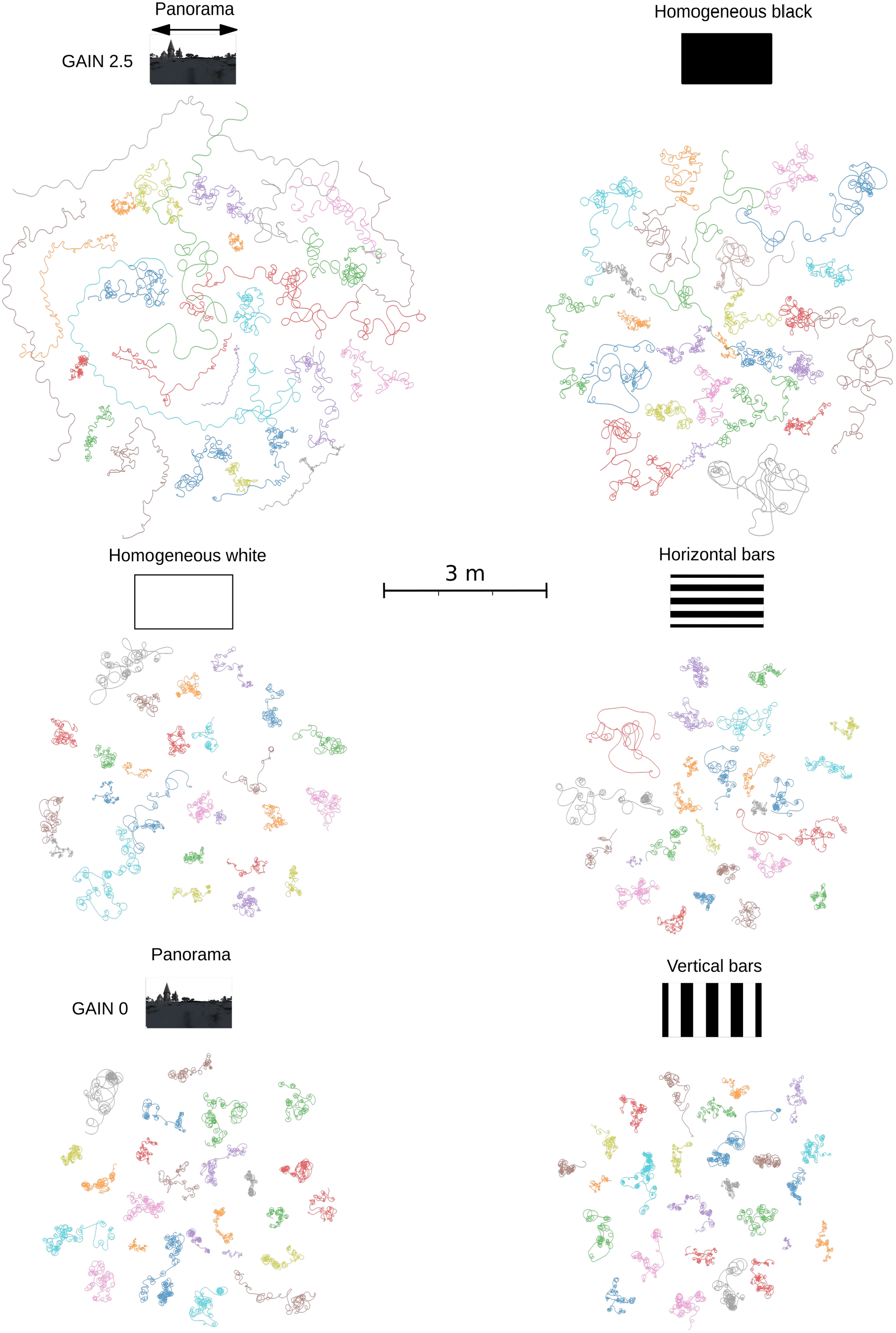
Reconstructed trajectories of all ants tested in the VR with different visual structures. Ants are getting stuck turning on the same side a lot when they are exposed to static vertical and horizontal bars, homogeneous white or a static panorama creating loops on their trajectories. Those loops seem slightly less tight when they are exposed to static horizontal bars or homogeneous white since they turn slower (Fig.3A).

## SUPPLEMENTARY INFORMATION 3

**Figure S3:**
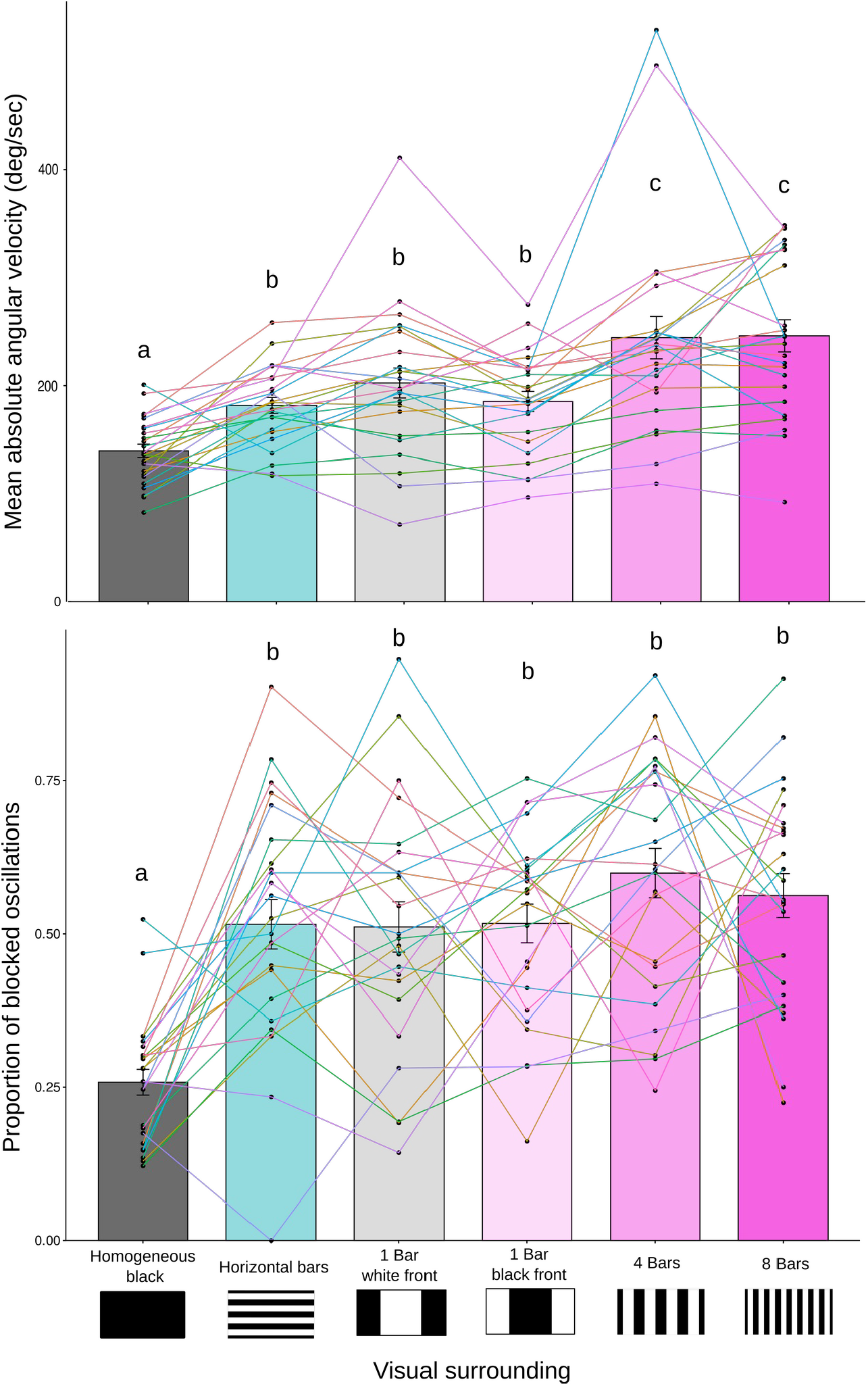
There is an effect of the number of vertical edges on the average angular speed of ants’ oscillations. Effect of the visual structure ants were exposed to in gain 0 on **(A)** the mean absolute angular velocity **(B)** and proportion of ants’ oscillations blocked above or under 0 deg/sec (without changing direction).Data based on 24 ants tested in the VR. The error bars shown in the bar plots represent the standard error of the means. Each point corresponds to the response of an individual ant while the lines are connecting the responses of the same ant across the different conditions. We used a LMM to test the effect of the visual structure ants were exposed to in gain 0 on the absolute angular velocities using the log transformation, and a GLMM for proportional data with the binomial distribution and the link function logit to test this effect on the proportion of blocked oscillations. For both models the identity of the ant was added as a random factor to account for repeated measurements and an observation level random factor was added to account for overdispersion in the GLMM. There was a significant effect of the visual structure both on the mean absolute angular velocities (LMM : 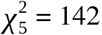, *P* < 0.001) and the proportion of blocked oscillations (GLMM : 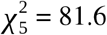, *P* < 0.001). Different letters correspond to significant post-hoc comparisons using the sequential Bonferroni correction after Holm.

## SUPPLEMENTARY INFORMATION 4

**Figure S4:**
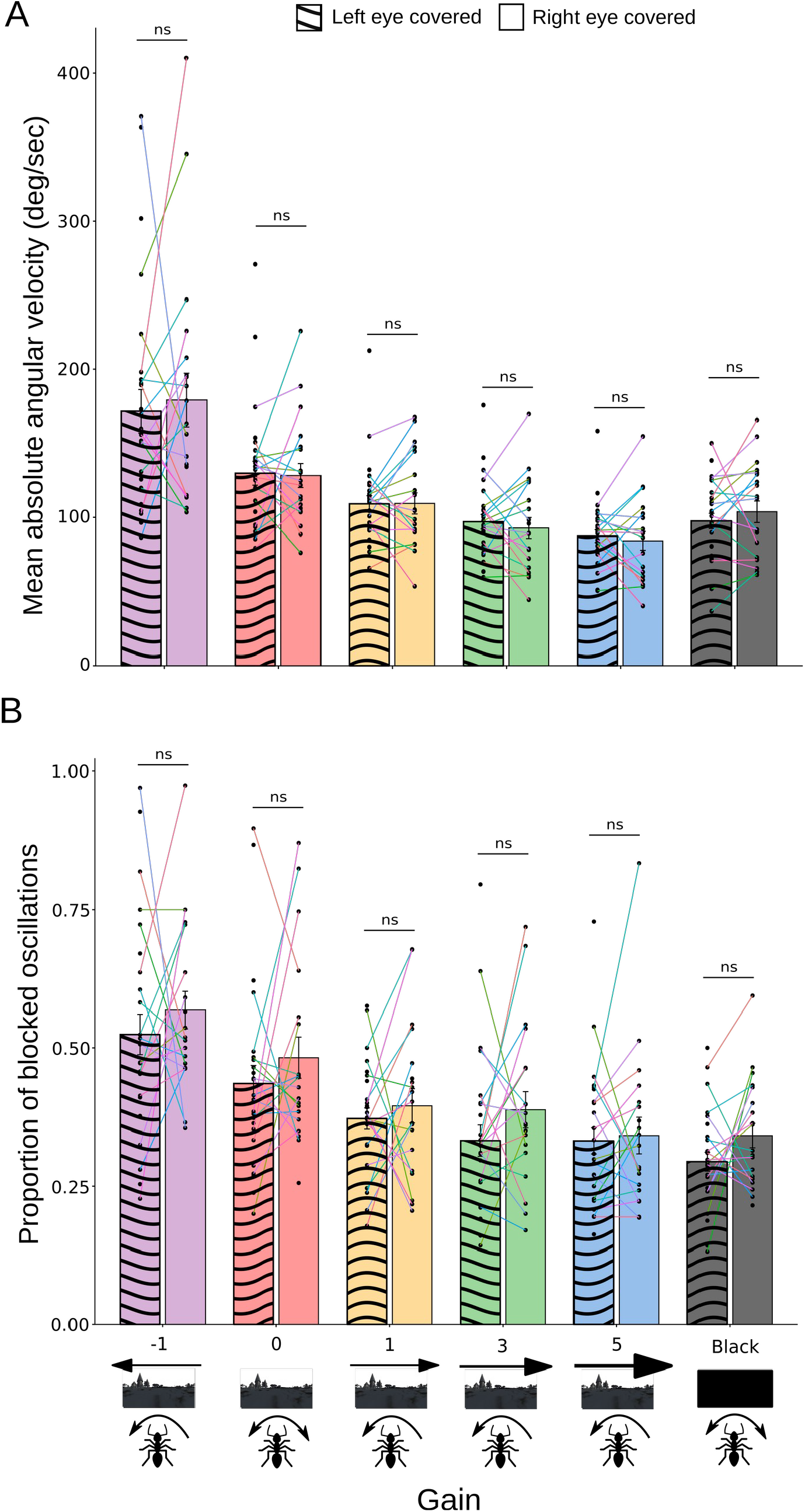
The side of the eye covered has no impact on the average angular speed or the proportion of blocked oscillation of ants. Effect of the side of the eye covered depending on the gain on **(A)** the mean absolute angular velocity **(B)** and proportion of ants’ oscillations blocked above or under 0 deg/sec (without changing direction). Data based on 27 ants tested in the VR including 20 ants tested with the right eye covered and 24 tested with the left eye covered. The error bars shown in the bar plots represent the standard error of the means. Each point corresponds to the response of an individual ant while the lines are connecting the responses of the same ant across the different conditions. We used a LMM to test for the effect of the side of the eye covered (factor with 2 levels: right or left eye covered) and the gain (factor with 6 levels: -1, 0, 1, 3, 5 and black) as well as the interaction between both on the absolute angular velocities using the log transformation, and a GLMM for proportional data with the binomial distribution and the link function logit to test for the effect of the side of the eye covered and the gain as well as the interaction between both on the proportion of blocked oscillations. For both models the identity of the ant and the trial was added as a random factor to account for repeated measurements and an observation level random factor was added to account for overdispersion in the GLMM. For both models only the gain had a significant effect (angular velocities, LMM : 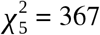, *P* < 0.001; blocked oscillation GLMM : 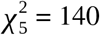, *P* < 0.001), both the interaction (angular velocities, LMM : 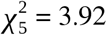, *P* = 0.562; blocked oscillation GLMM : 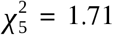, *P* = 0.888) and the effect of the side of the eye covered (angular velocities, LMM : 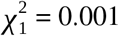, *P* = 0.973; blocked oscillation GLMM : 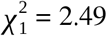, *P* = 0.115) were not significant.

